# Scaling recipes for single-cell RNA sequencing foundation models: when do scaling laws hold?

**DOI:** 10.64898/2026.08.31.747783

**Authors:** Federico Borra, Giacomo Cirò, Arianna Castellini, Giovanni Gatti, Andrea Tangherloni, Francesca M. Buffa

## Abstract

Deep learning models exhibit empirical scaling laws whereby performance changes predictably with model size, dataset size, and training compute. Although these relationships are well established in domains such as language and image modelling, their applicability to biological data remains unclear. Here, we investigate scaling behaviour in foundation models trained on large collections of single-cell transcriptomes. We show that pre-training loss decreases systematically with model capacity and training compute, exhibiting a power-law dependence on model size. The strength and regularity of these trends differ between model formulations. We identify and quantify empirical relationships linking the optimal learning rate and depth-to-width ratio to model size and depth or compute. These results demonstrate that scaling principles extend to transcriptomic modelling. More broadly, they provide a quantitative framework for estimating the expected returns from additional resources and selecting suitable hyperparameters and architectures, thereby supporting the development of increasingly capable foundation models for omics data.

## 1 Main

**Fig. 1:**
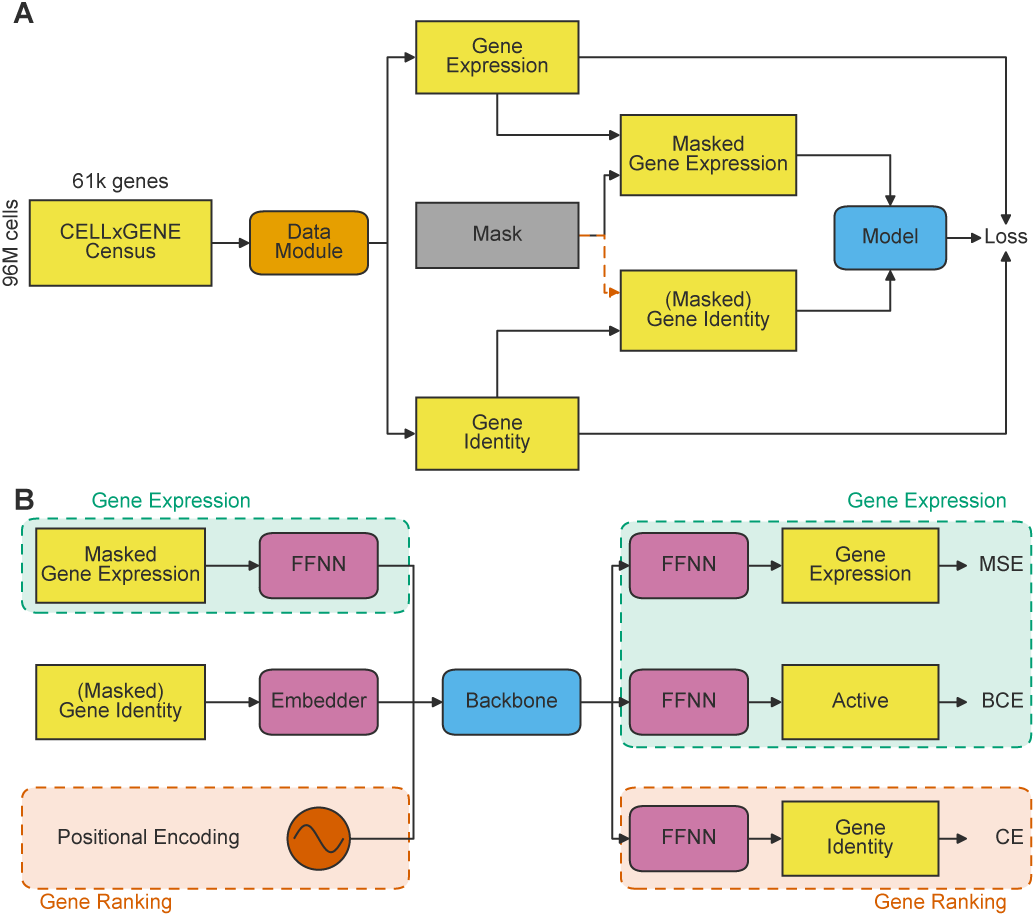
Modular data pipeline and model formulations. **(A)** CELLxGENE Census data comprising 96,591,226 cells and 61,497 RNA features are converted into paired gene-expression values and gene identifiers. The data module performs formulation-specific gene selection and masking, while retaining the original values or identifiers as prediction targets. **(B)** The two formulations use a common transformer-encoder architecture but distinct input representations and prediction heads. In the Binned Gene Expression formulation (green), gene identifiers are embedded, and masked expression bins are projected through a feed-forward neural network (FFNN); separate heads predict expression bins using mean squared error (MSE) and gene activity using binary cross-entropy (BCE). In the Ranked Gene Identity formulation (red), gene embeddings are combined with positional encodings representing expression rank, and a gene-identity head reconstructs masked gene identifiers using cross-entropy (CE). The two formulations are trained separately.

The recent success of deep learning across diverse domains, most notably natural language processing and image generation, has been remarkable. Despite these advances, our theoretical understanding of why large models perform so well remains incomplete. Nonetheless, consistent empirical regularities have emerged, enabling quantitative prediction of their performance [1–3]. These scaling laws describe how model performance varies as a power-law function of key resources, including dataset size, model size, and training compute, provided that no other factor becomes a limiting bottleneck.

At the same time, advances in single-cell sequencing have transformed the study of cellular biology. Single-cell RNA sequencing (scRNA-seq) enables detailed characterisation of cellular heterogeneity, lineage relationships, and disease-associated transcriptional programmes. Large collaborative initiatives, such as the Human Cell Atlas [4], have produced transcriptomic atlases containing tens of millions of cells, while continued technological advances are broadening the range of molecular modalities that can be measured. These rapidly expanding datasets offer unprecedented opportunities to investigate cellular systems at scale, while also presenting substantial computational challenges.

Foundation models pre-trained on large-scale datasets have recently emerged as a promising means of harnessing the growing volume of biological data [5, 6]. Across other domains, transformer-based models have demonstrated an ability to learn general-purpose representations that can be adapted to a wide range of downstream tasks [7]. Building on this paradigm, several foundation models for transcriptomic data have recently been introduced, including Geneformer and scGPT, which learn representations of genes and cells from extensive collections of single-cell expression profiles.

Very recently, there has been a shift towards characterising these scaling laws in single-cell foundation models [8–15]. Existing analyses, however, remain partial: they are often qualitative, focus on a single model architecture, and primarily examine scaling with computational resources, with limited consideration as to how scaling behaviour can inform hyperparameter selection. In this work, we take a step towards a more comprehensive examination of the scaling behaviour of transcriptomics foundation models. We develop a standardised pipeline that enables controlled and fair comparisons across multiple experimental axes. Its modular design allows preprocessing strategies, pre-training objectives, loss functions, and model architectures to be varied independently, while holding other conditions fixed, including the underlying data and the randomised order in which samples are presented to the models.

Finally, we assess whether improvements in pre-training loss translate into better downstream performance, considering cell-type classification, batch correction, and, more broadly, the quality of the learned representations as measured by their clustering properties. Through quantitative analysis, we empirically demonstrate that scaling behaviour occurs in single-cell foundation models and characterise it using scaling coefficients. We further show that similar regularities arise not only under resource scaling (e.g., increasing the number of parameters or the amount of compute), but also when varying other hyperparameters and architectural properties, including the learning rate and depth-to-width ratio. Although scaling behaviour has previously been demonstrated in large language models, we find that some scaling coefficients differ in magnitude from those reported for language models and that, in one case, the sign of the coefficient is reversed.

These findings highlight the importance of comprehensively characterising a model’s scaling behaviour before committing to large-scale training, particularly when scaling laws are used to guide architectural decisions and optimise the resources required to achieve a target level of performance.

## 2 Results

### 2.1 Scaling Laws at Different Context Lengths

We trained a broad range of models on the CELLxGENE Census [16] using two model formulations: (*i*) Ranked Gene Identity prediction, following a Geneformer-like approach and optimised using masked-token cross-entropy; and (*ii*) Binned Gene Expression prediction, following an scGPT-like approach and jointly optimised using expression MSE and gene-activity BCE.

Figure 2 shows that training loss decreased across the tested batch sizes and model widths for both formulations, indicating stable optimisation throughout the explored range. At a fixed optimiser step, loss was generally lower for larger batches and wider models. However, because larger batches process more cells and incur more training compute per step, these comparisons do not isolate the effect of gradient noise or identify a critical batch size.

**Fig. 2:**
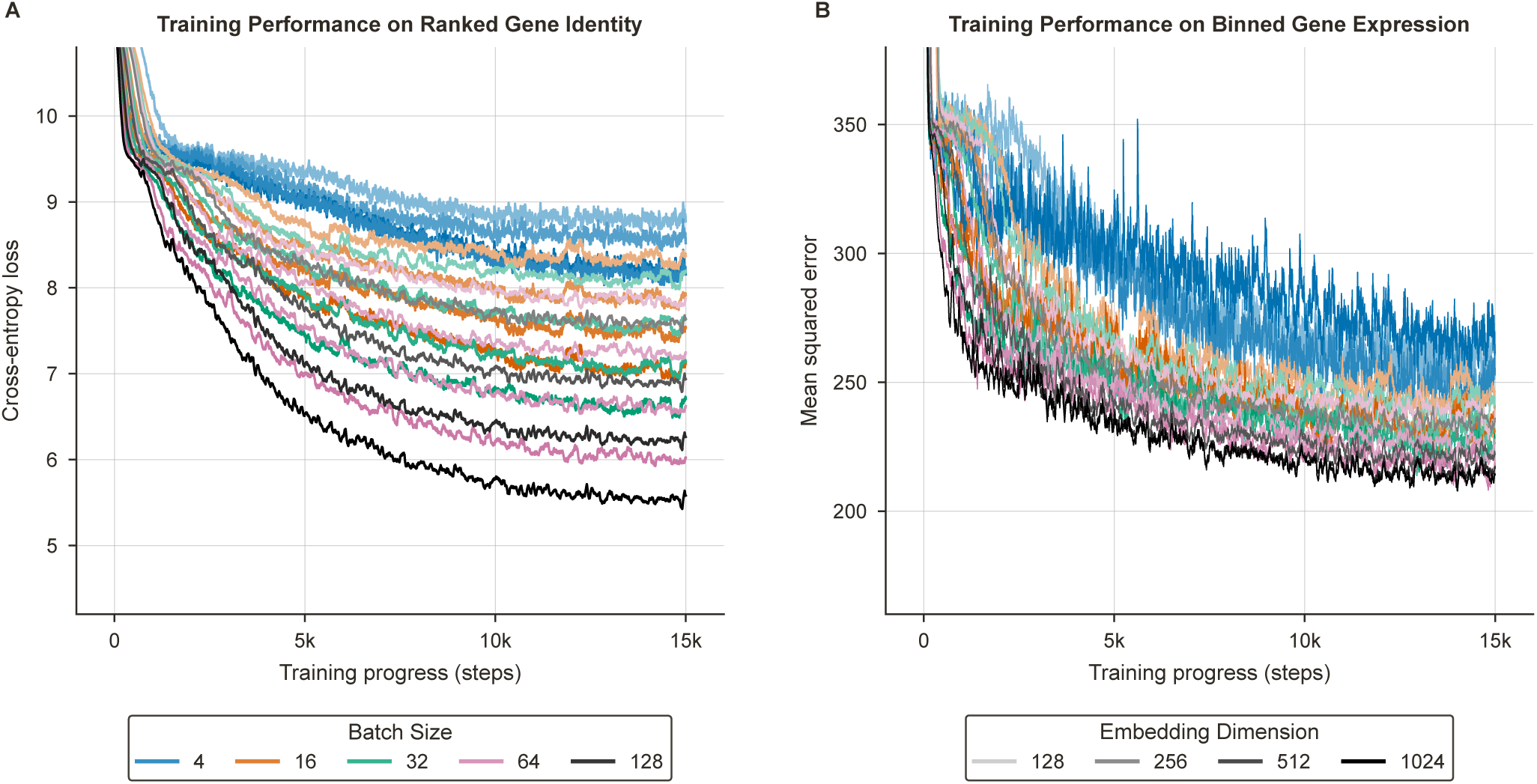
Training loss dynamics across batch sizes. Exponentially smoothed training loss (α = 0.05) over 15,000 optimiser steps for **(A)** the Ranked Gene Identity formulation, evaluated using masked-token cross-entropy, and **(B)** the Binned Gene Expression formulation, evaluated using expression-head MSE on masked positions with non-zero targets. All models used a context length of 256, six transformer blocks, four attention heads, embedding dimensions of 128, 256, 512, or 1,024, and physical and effective batch sizes of 4, 16, 32, 64, or 128, without gradient accumulation. Colour denotes batch size, and shade denotes embedding dimension; lower values indicate better training performance.

We selected a batch size of 32 for the subsequent experiments as a practical compromise between per-step memory requirements and the number of cells processed per update, and held it fixed throughout the principal sweeps.

Figure 3A shows that the rank-based model exhibits scaling behaviour at every context length tested. Cross-entropy loss decreases smoothly with the number of non-embedding parameters, producing an approximately linear relationship on log–log axes. In addition, the fitted scaling exponent increases monotonically with context length, from k = 0.022 at a context length of 128 to k = 0.092 at a context length of 2,048. This indicates that additional model capacity becomes increasingly beneficial as more genes are included in the input context.

**Fig. 3:**
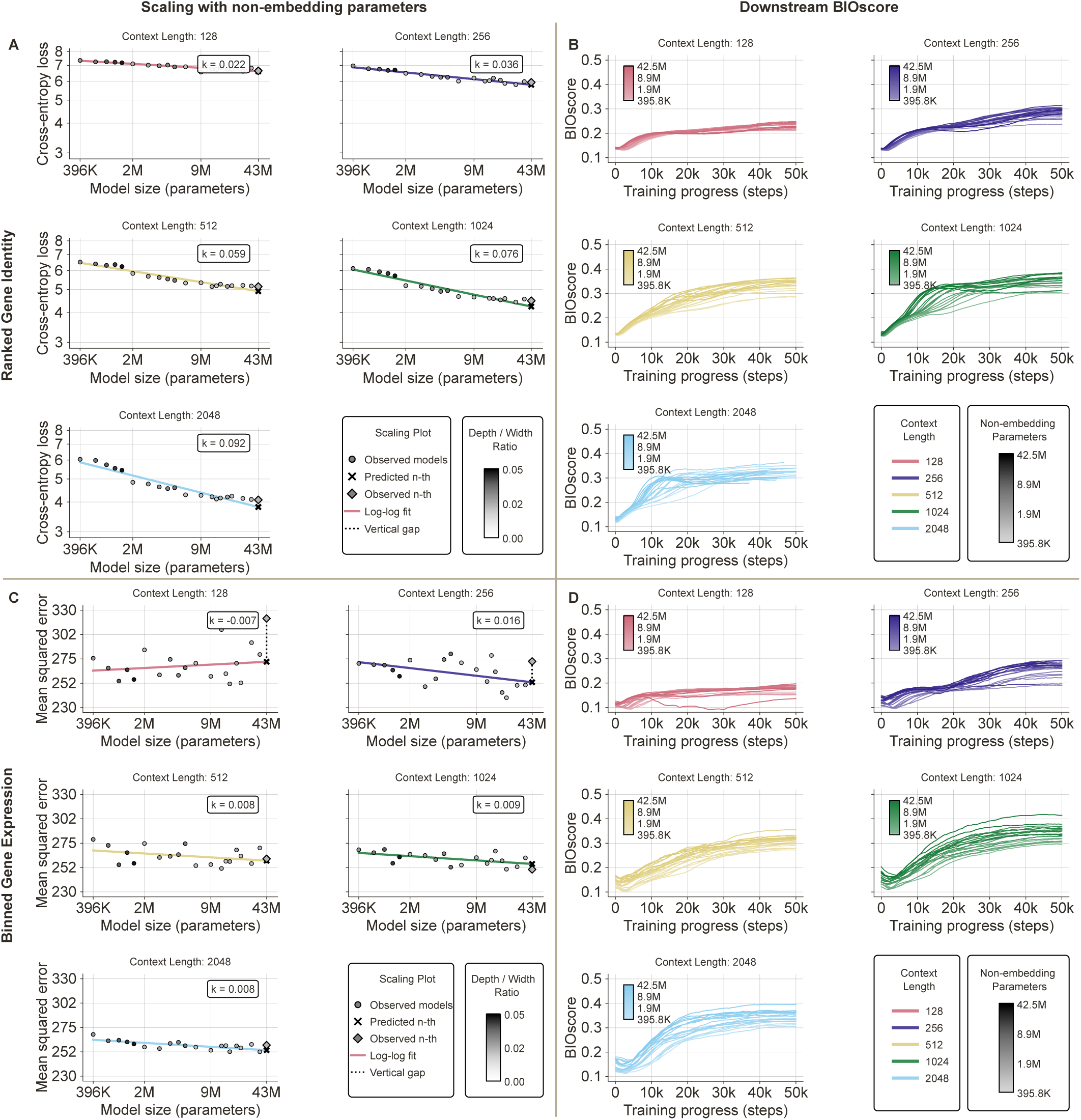
Scaling of pre-training loss and downstream BIOscore with model size and context length. **(A, C)** Pre-training loss as a function of non-embedding parameters, N, for Ranked Gene Identity prediction measured by cross-entropy **(A)** and Binned Gene Expression prediction measured by MSE **(C)**. Each subplot corresponds to a different context length (128-2,048), and marker shading indicates the depth-to-width ratio. Solid lines show log-log fits of the scaling law L ∝ N *^−k^*. Crosses and diamonds denote the predicted and observed losses, respectively, for the target model, while dotted lines indicate the prediction gap. **(B, D)** Downstream BIOscore trajectories over 50,000 training steps for the Ranked Gene Identity **(B)** and Binned Gene Expression **(D)**. Colour denotes context length, and darker shades indicate higher non-embedding parameter count. Ranked Gene Identity exhibits stronger loss scaling than Binned Gene Expression, while BIOscore trajectories show how these pre-training trends translate into the quality of learned representations.

This result is intuitive: longer contexts provide more information for the model to learn complex relationships among genes. In the limit case of a context length of one, the model has no neighbouring genes from which to infer higher-order relationships. Once the marginal gene statistics have been captured, additional model capacity should therefore provide little benefit, and scaling is expected to be negligible.

By contrast, the Binned Gene Expression model exhibits substantially weaker and less consistent scaling behaviour (Figure 3C). Although the mean squared error (MSE) generally decreases with model size, the fitted coefficients are small and do not increase monotonically with context length. Instead, the principal effect of increasing the context length appears to be a reduction in the dispersion of the observed losses around the fitted power-law trajectory.

Because the two formulations use different loss functions (i.e., cross-entropy for Ranked Gene Identity prediction and MSE for Binned Gene Expression prediction), the absolute loss values are not directly comparable. We therefore focus on the fitted scaling coefficients and the consistency of the observed scaling trends rather than on the magnitudes of the losses.

#### 2.1.1 Downstream Performance

To assess whether reductions in pre-training loss translate into improved downstream performance, Figures 3B and 3D report the results of our evaluation procedure. We summarise performance using BIOscore, a custom composite metric that combines the classification accuracy of a ridge-regularised linear probe on coarse cell-type labels with several clustering-quality metrics computed from the cell embeddings using both the fine-grained labels provided by CELLxGENE and a coarse-grained version derived from them (Sections 4.2.1 and 4.10). The individual BIOscore components exhibited broadly concordant trends across training and model configurations. We therefore aggregated them into a single composite score to provide a compact visualisation, while reporting the component metrics separately in the Supplementary Information. These results provide indicative, rather than definitive, evidence of an association between pre-training and downstream performance. Previous work has shown, for instance, that single-cell foundation models do not consistently outperform simpler baselines [15, 17, 18]. It is unclear whether this is due to task saturation, ill-defined or noisy objectives, or limitations of the foundation models themselves. With this experiment, however, we are interested in the trend of improvement in the downstream tasks rather than in absolute performance. As training progresses and pre-training loss decreases, the BIOscore generally increases. The learned representations also tend to improve with model size and context length.

These trends should be interpreted with caution, as several confounding factors remain. In particular, embedding dimensionality increases with model width, with immediate and sizeable repercussions for the linear probe’s expressive power. There is, in fact, a linear relationship between the number of parameters in the linear probe and the embedding dimensionality. This gives wider models an advantage under this evaluation protocol. As a consequence, part of the apparent improvement with model size may reflect differences in embedding dimensionality rather than representational quality alone.

Finally, the two model families reach comparable BIOscores, particularly at longer context lengths. Within the range of model sizes and compute budgets considered here, neither formulation therefore exhibits a consistent advantage in downstream performance. This conclusion is limited to the scales and evaluation procedure examined in this study.

### 2.2 Optimal Learning Rate Scaling

We perform an extensive sweep to determine the optimal learning rate across combinations of parameter count and model depth for both the Ranked Gene Identity and Binned Gene Expression formulations. Figures 4A and 4D show that, at any fixed depth, the optimal learning rate decreases with the number of non-embedding parameters according to an approximate power law.

**Fig. 4:**
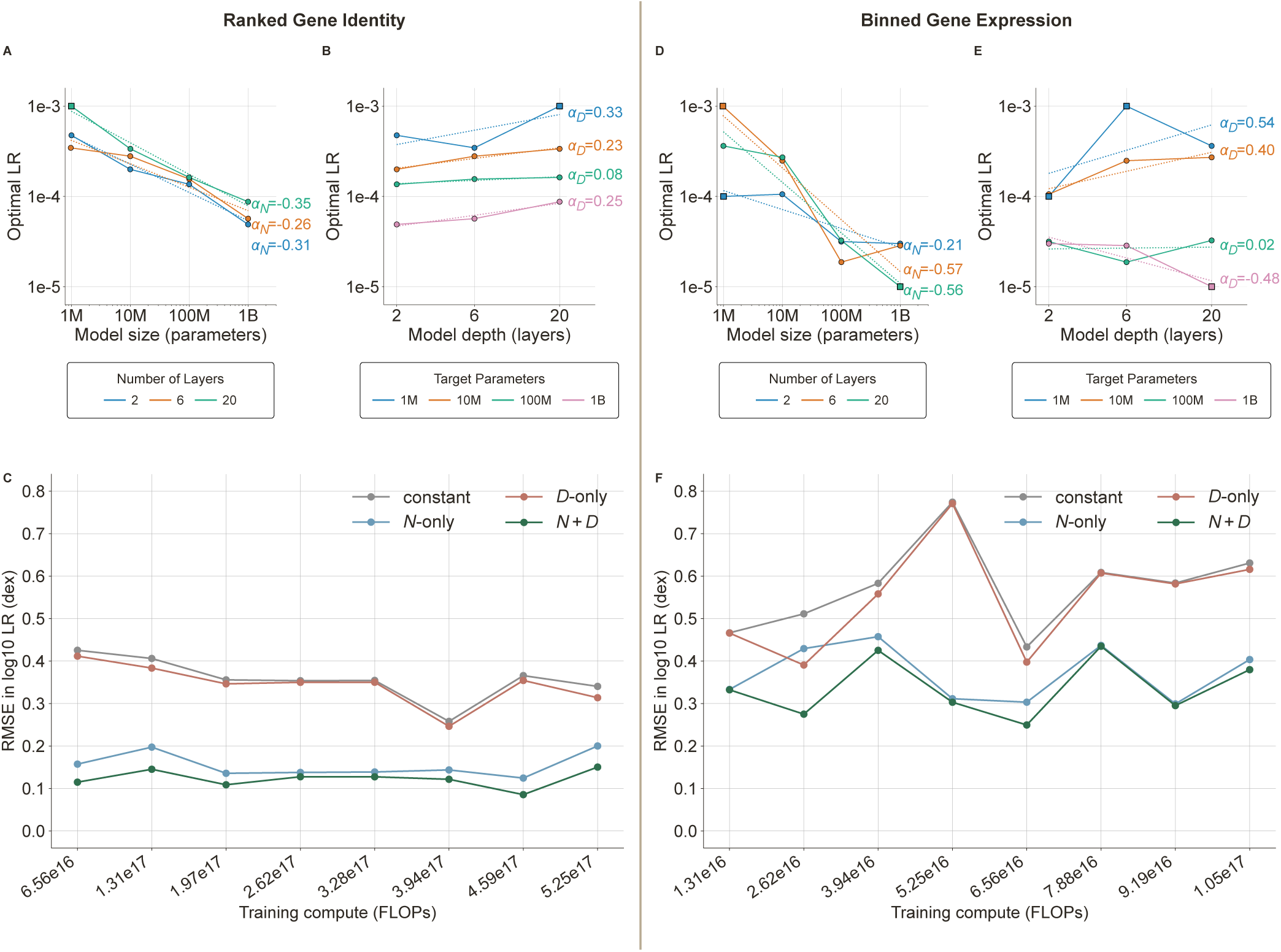
Scaling of the optimal learning rate with model size, depth, and training compute. Results are shown for the Ranked Gene Identity formulation (A–C) and the Binned Gene Expression formulation (D–F). **(A, D)** Optimal learning rate as a function of the number of non-embedding parameters at the penultimate compute slice, corresponding to 65.625% of the total training budget (4.59 × 10^17^ FLOPs for Ranked Gene Identity prediction and 9.19 × 10^16^ FLOPs for Binned Gene Expression prediction). Colours denote model depth. **(B, E)** Optimal learning rate as a function of depth for models matched to non-embedding-parameter targets ranging from 1 million to 1 billion. Colours denote the target parameter count. Solid lines connect the estimated optima, dotted lines show the corresponding power-law fits, and the fitted size and depth exponents, α*_N_* and α*_D_*, are reported beside the corresponding curves. Circles indicate optima estimated by quadratic approximation, whereas squares identify sampled optima retained because the winner was at a search boundary or did not have a complete eligible three-point fitting window. **(C, F)** Predictive accuracy of four learning-rate scaling rules across compute slices: a constant learning rate, scaling with non-embedding parameter count only (N-only), scaling with depth only (D-only), and joint scaling with both non-embedding parameter count and depth (N +D). Error is measured as the root-mean-square error of the predicted log_10_ learning rate in dex; lower values indicate better predictions. Across both formulations, the joint N +D model generally provides the most accurate description, indicating that optimal learning-rate selection depends on both model size and architectural shape.

At the shown compute slice, the magnitudes of the fitted size exponents, |α*_N_* |, range from 0.26 to 0.35 for the Ranked Gene Identity model and from 0.21 to 0.57 for the Binned Gene Expression model. These values are substantially larger than the exponent magnitude of approximately 0.14 reported in comparable experiments on transformer language models [2]. This discrepancy highlights the importance of repeating such empirical analyses in new modelling settings rather than assuming that an established scaling recipe will transfer across domains.

Looking at the same results by model depth, while holding the parameter count approximately constant, reveals a different trend (Figures 4B and 4E). For the Ranked Gene Identity model, the optimal learning rate generally increases with depth, with positive fitted depth exponents, α*_D_*, ranging from 0.08 to 0.33. This direction is the reverse of that reported for language models, where the optimal learning rate decreases with depth [19, 20]. For the Binned Gene Expression model, however, the dependence is less consistent and varies with the target parameter count. Specifically, α*_D_* ranges from −0.48 to 0.54, and the optimal learning rate decreases with depth at the largest parameter target. These results indicate that parameter count alone is insufficient to determine the optimal learning rate. Indeed, both model depth and model formulation must be considered.

A joint power law incorporating both model depth and parameter count provides a more accurate characterisation of optimal learning-rate scaling than either variable alone, as shown in Figures 4C and 4F. We therefore use this joint relationship in the analyses that follow.

The difference in fit quality between the two formulations is nevertheless substantial. At the highlighted compute slice, the joint power law achieves an RMSE of approximately 0.09 dex for the Ranked Gene Identity model, compared with approximately 0.30 dex for the Binned Gene Expression model.

Since the error is measured on a logarithmic scale, this corresponds to a substantially smaller deviation for the Ranked Gene Identity model than for the Binned Gene Expression model. An RMSE of 0.09 dex corresponds to a characteristic multiplicative factor of 10^0.09^ ≈ 1.23, equivalent to deviations of approximately +23% or -19% from the optimum. By comparison, an RMSE of 0.30 dex corresponds to a factor of 10^0.30^ ≈ 2.00, or approximately +100% and -50%.

This higher residual error provides further evidence that the Binned Gene Expression model exhibits less regular scaling behaviour than the Ranked Gene Identity model, consistent with the trends observed in Figure 3.

### 2.3 Depth Scaling with Compute Budget

Having established a scaling law for the optimal learning rate, we use it to investigate how optimal model depth varies with model size. We evaluate a denser grid of depths at approximately logarithmically spaced non-embedding parameter targets ranging from 1 million to 1 billion. For each configuration, we use the learning rate predicted by the joint scaling law derived in the previous section.

We fit a quadratic response surface that models pre-training loss jointly as a function of parameter count, training compute, and depth-to-width ratio (as detailed in Methods; Section 4.9, extending prior approaches in neural scaling literature such as quadratic isoFLOP profiles or multidimensional loss surfaces [3, 21]).

For the Ranked Gene Identity model, the fitted surface indicates that the optimal depth-to-width ratio decreases as model size increases at fixed compute, favouring relatively wider models. Conversely, at a fixed parameter count, the optimal ratio increases with the compute budget, favouring relatively deeper models. Within the scaling regime covered by our experiments, this surface therefore provides an empirical rule for selecting model depth and width for a given parameter count and compute budget to minimise the predicted loss.

For the Binned Gene Expression model, however, the full quadratic response surface provides a substantially poorer description of the data. First, its held-out predictions exhibit greater dispersion (Figure 5E) than those of the Ranked Gene Identity model (Figure 5B). The R^2^ values decrease from 0.985 to 0.876 under random five-fold cross-validation and from 0.983 to 0.868 under leave-one-compute-slice-out validation. More importantly, the optimum inferred for the Binned Gene Expression model is highly unstable under bootstrap resampling. Comparing Figures 5C and 5F, the Ranked Gene Identity model produces narrow and consistent estimates of the optimal depth-to-width ratio, whereas the estimates for the Binned Gene Expression model span as many as four orders of magnitude. Moreover, approximately 15% of the bootstrap-fitted surfaces are not convex and therefore do not yield a stationary point corresponding to a minimum. Consequently, inferring the optimal depth-to-width ratio from the fitted surface is justified for the Ranked Gene Identity model but not for the Binned Gene Expression model.

**Fig. 5:**
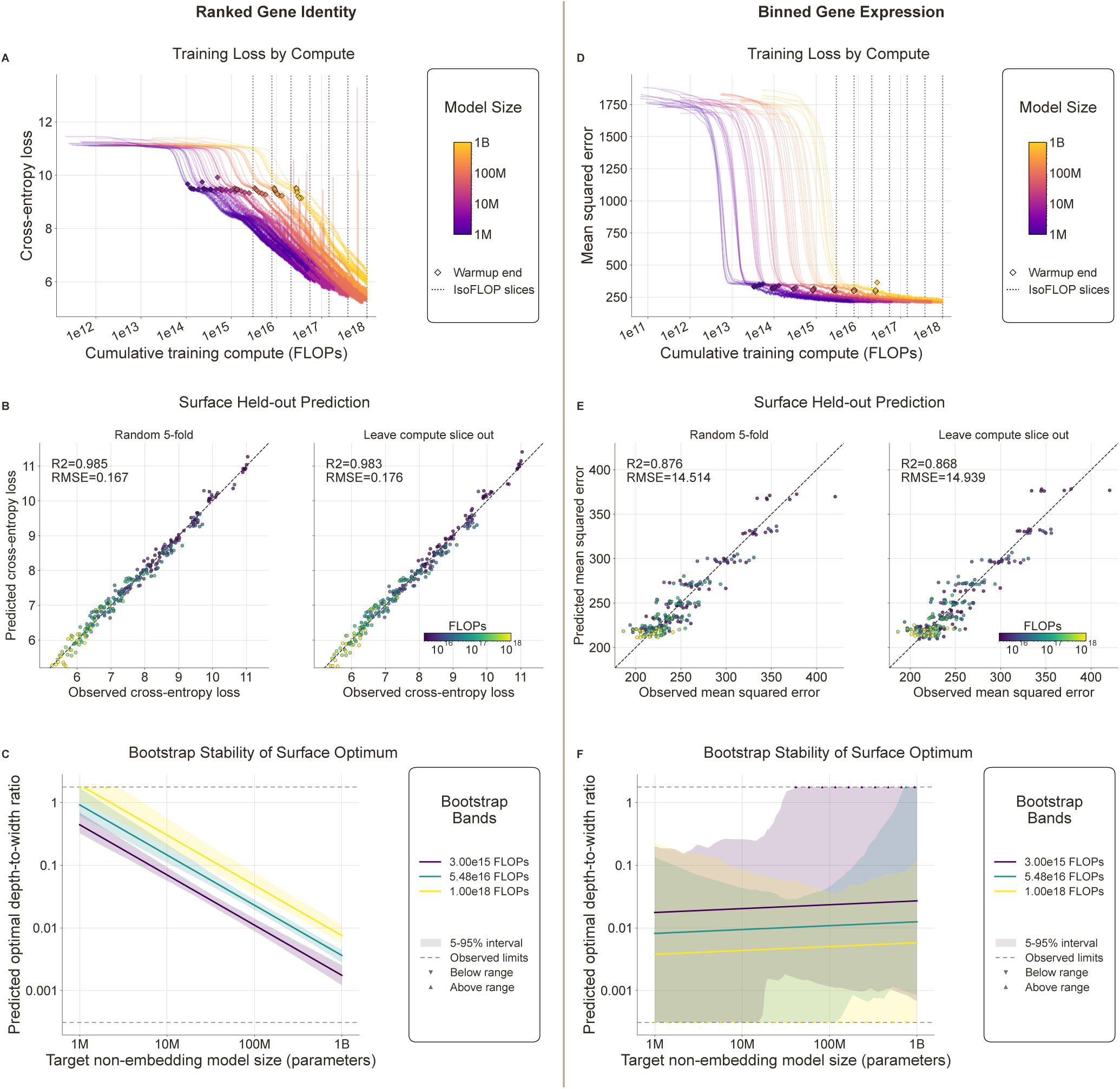
Loss-surface modelling and estimation of compute-optimal model shape. Results are shown for the Ranked Gene Identity formulation (A–C) and the Binned Gene Expression formulation (D–F). **(A, D)** Training loss as a function of cumulative compute for models ranging from 1 million to 1 billion non-embedding parameters. Colour denotes model size, diamonds indicate the end of learning-rate warm-up, and vertical dotted lines mark the isoFLOP slices used to construct the loss surfaces. **(B, E)** Held-out predictive performance of the fitted loss surfaces under random five-fold cross-validation and leave-one-compute-slice-out validation. Each point compares predicted and observed loss and is coloured by training compute; dashed lines indicate perfect agreement. The corresponding R^2^ and root MSE (RMSE) values are reported in each panel. **(C, F)** Predicted optimal depth-to-width ratio as a function of the target non-embedding parameter count at three compute budgets. Solid lines show the estimated optima, and shaded regions represent the 5th–95th percentile intervals obtained by bootstrap resampling. Horizontal dotted lines delimit the range of architectural ratios represented in the experiments, while triangles identify predictions below or above this observed range. The Ranked Gene Identity loss surface yields accurate held-out predictions and stable architectural optima, whereas the Binned Gene Expression surface exhibits lower predictive accuracy and substantially greater uncertainty, indicating that its optimal model shape is less reliably identifiable.

These results emphasise the empirical nature of scaling behaviour. A model formulation may offer an important practical advantage not only through its absolute performance, but also through the regularity and predictability of its scaling relationships, which enable more reliable hyperparameter selection and extrapolation.

Figure 6 shows contours of the fitted quadratic response surface for the Ranked Gene Identity model. We omit the corresponding surface for the Binned Gene Expression model because its instability makes the inferred optima unreliable. The predicted optimal depth-to-width ratio is overlaid on each surface slice, together with the observed configurations used to fit the surface where applicable.

**Fig. 6:**
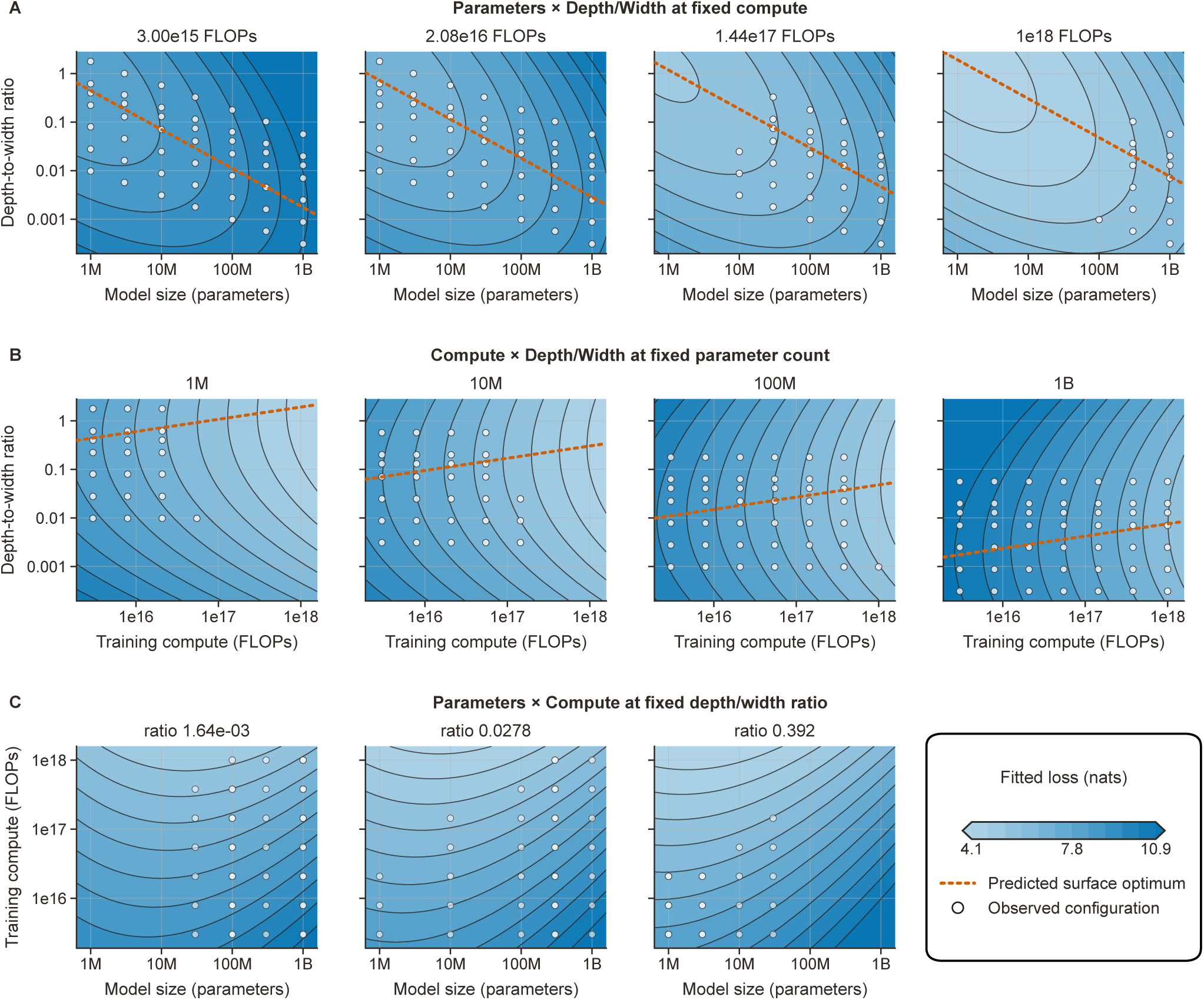
Two-dimensional slices through the quadratic loss surface fitted for the Ranked Gene Identity formulation. The surface models pre-training loss jointly as a function of non-embedding parameter count, training compute, and depth-to-width ratio. **(A)** Fitted loss as a function of non-embedding parameter count and depth-to-width ratio at four fixed compute budgets, ranging from 3.00 × 10^15^ to 1.00 × 10^18^ FLOPs. **(B)** Fitted loss as a function of training compute and depth-to-width ratio at fixed non-embedding-parameter targets of 1 million, 10 million, 100 million, and 1 billion. Orange dashed lines trace the depth-to-width ratio that minimises the fitted loss at each parameter count or compute budget. **(C)** Fitted loss as a function of non-embedding parameter count and training compute at three fixed depth-to-width ratios. White circles denote observed training configurations, contour lines connect configurations with equal predicted loss, and colour indicates the fitted loss, with lighter shades corresponding to lower values. All axes are logarithmic. The fitted surface predicts that the optimal depth-to-width ratio decreases with parameter count at fixed compute but increases with the available compute at fixed parameter count, favouring relatively wider models as model size grows and relatively deeper models as the training budget increases.

## 3 Discussion

Our results show that foundation models trained on single-cell transcriptomic data can exhibit scaling behaviour similar to that observed in language, vision, and audio models. In particular, the Ranked Gene Identity model displays a predictable improvement in pre-training loss as model size and compute increase. This regularity is encouraging because, within the regime examined, it enables estimation of the expected benefit of additional resources before committing to large-scale training. However, the weaker, less stable scaling observed in the Binned Gene Expression model suggests that this behaviour cannot be assumed to hold equally across model formulations.

The regularities identified here extend beyond resource variables (e.g., parameter count and training compute) to include optimisation hyperparameters and architectural choices. In particular, the optimal learning rate depends systematically on model size and depth, while the fitted response surface for the Ranked Gene Identity model relates the optimal depth-to-width ratio to both parameter count and compute. Such relationships are especially valuable at large scales, where exhaustive trial-and-error optimisation becomes prohibitively expensive. Conversely, using poorly tuned hyperparameters can waste substantial computational resources and yield suboptimal performance despite an otherwise sufficient training budget.

The workflow introduced in this study also provides a practical means of reducing the complexity of hyperparameter optimisation. Rather than evaluating every combination in an exhaustive Cartesian grid, the search can be decomposed into lower-dimensional sweeps whose results are subsequently combined through joint scaling laws and response surfaces. This factorised approach replaces a search whose cost grows multiplicatively across hyperparameter axes with a sequence of more tractable empirical estimation problems.

Although we demonstrate these properties using models of static single-cell transcriptomic states, the same methodology could be particularly valuable for perturbation-response models. If similar scaling regularities hold, empirical scaling laws could help determine suitable architectures, hyperparameters, and training budgets for increasingly ambitious models of cellular behaviour. Such an application requires at least two conditions: (*i*) the training objective must align closely with the biological behaviour of interest, and (*ii*) the task must be sufficiently complex and informative to benefit from substantial model capacity. Under these conditions, the approach developed here could contribute to the design of future “virtual cell” models. Analogous procedures may also apply to other data-intensive areas of biomedicine.

Potential examples include models of histopathological images, spatial omics data, biomolecular structures, and multimodal clinical records. These settings often require substantial model capacity and computational investment, making reliable estimates of scaling behaviour and optimal hyperparameters especially valuable.

An important open question is why Ranked Gene Identity prediction exhibits substantially more regular scaling than Binned Gene Expression prediction. A controlled interpolation between the two representations, for example, by progressively refining the expression discretisation towards a rank-based representation, could help determine whether the difference arises primarily from the input representation, the prediction objective, or their interaction.

The principal limitations of this approach are its empirical and setting-specific nature. The fitted relationships provide no guarantee that observed trends will continue arbitrarily far beyond the range of model sizes and compute budgets examined. Moreover, the scaling coefficients depend on the data, preprocessing procedure, pre-training objective, architecture, optimiser, and evaluation protocol. As demonstrated by the contrasting behaviour of the two models studied here, some formulations are considerably more amenable to scaling analysis than others, and the resulting laws are unlikely to transfer unchanged to new configurations. Scaling laws should therefore be treated as locally validated empirical models and reassessed at intermediate scales before they are used to guide substantially larger training runs. Future work could also investigate whether combining this empirical framework with parameterisation-based hyperparameter-transfer methods improves its reliability and portability [19, 20, 22, 23].

## 4 Methods

### 4.1 Study design

We developed a common training and evaluation pipeline to study empirical scaling relationships in transformer encoders trained on scRNA-seq profiles. The pipeline held the data source, cell split, sampling procedure, optimiser family, and evaluation protocol fixed while varying the input representation, pre-training objective, context length, encoder width, encoder depth, learning rate, and training budget. We studied two formulations: *(i)* masked gene identity prediction from expression-ranked gene sequences (the Ranked Gene Identity formulation), and *(ii)* joint prediction of masked gene activity and binned expression (the Binned Gene Expression formulation). The two formulations shared the same encoder implementation but used representation-specific input modules and output heads.

Unless stated otherwise, each architecture was trained once. The absence of independent training-seed replicates means that the uncertainty estimates reported below quantify sensitivity to sampled architectures or surface observations, rather than variation across model initialisations.

### 4.2 Single-cell data and data splits

Experiments used the human RNA measurement from a local TileDB-SOMA copy of the 8 November 2025 CELLxGENE Census release (release identifier 2025-11-08). We retained observations that satisfied is primary data == True and read the normalised expression layer. This filter yielded 96,591,226 cells across 61,497 RNA features. No additional cell-level quality-control filter, gene filter, or selection of highly variable genes was applied. Gene tokens were zero-based indices into the complete Census RNA feature ordering, whose feature identifiers were Ensembl gene identifiers. Cells were divided once into training, validation, and test partitions with tiledbsoma ml.ExperimentDataset.random split and seed 42. Each was capped at 30,000 cells by reducing its fraction size as needed, with the remainder assigned to training. The resulting partitions contained 96,531,226 training cells, 30,000 validation cells, and 30,000 reserved test cells. The split was performed at the cell level without stratification or grouping by source dataset or donor so that a source dataset could contribute cells to multiple partitions. The test partition was reserved but not used in the analyses reported here. Training observations were shuffled by TileDB-SOMA using seed 42, an I/O batch size of 65,536 observations, and shuffle chunks of 64 observations. All model families used the same split and shuffle seed. Cells were sampled according to their natural frequency in the filtered Census, without reweighting or balancing by dataset, tissue, donor, or cell type. No process-wide random seed was set by the training entry point or cluster launch scripts. Consequently, PyTorch [24] model initialisation and dropout, together with NumPy-based [25] gene subsampling and masking, depended on the generator states initialised separately in each process and were not exactly reproducible from the experiment configuration alone.

Cell-type strings, Cell Ontology [26] term identifiers, and source-dataset identifiers were retained for downstream evaluation. The dataset of origin was used as the batch label, due to the absence of more granular information. Fine cell-type labels were the CELLxGENE annotations.

#### 4.2.1 Ontology-derived coarse cell-type labels

Coarse cell types were a study-specific reduction of the fine CELLxGENE annotations, rather than a field supplied by CELLxGENE. The mapping was constructed from the Cell Ontology term frequencies in the fixed 30,000-cell validation partition and was then held constant across model checkpoints and formulations. The validation partition contained 631 distinct input labels, including the special label unknown. We parsed the 17 March 2026 release of the Cell Ontology (cl.owl; ontology version 2026-03-17) and retained explicit Cell Ontology to Cell Ontology rdfs:subClassOf relations. Multiple-parent relations were retained, so the resulting structure was treated as a directed acyclic graph rather than as a single-parent tree.

We reduced the observed ontology terms to G = 30 anchors with a deterministic, frequency-adaptive bottom-up procedure, where G denotes the target number of coarse groups. Labels equal to unknown, na, or missing were omitted when fitting the reduction. Initially, every observed fine term was its own anchor. For each term, ontology depth was defined as the minimum number of subclass edges from any ontology root. Two counts were then maintained: a local count, equal to the number of validation cells currently assigned to an anchor, and a global propagated count, obtained by adding each fine term’s count to that term and to each of its unique ancestors.

While more than 30 anchors remained, the movable anchor with the smallest local cell count was selected for ascent. Ties were resolved first in favour of the deeper term and then by lexicographic Cell Ontology identifier. If the selected term had multiple direct parents, its parent was selected by the following ordered criteria: (i) a parent already present among the current anchors; (ii) a parent that was an ancestor of another current anchor; or (iii) any remaining direct parent. Within each class, the parent with the largest global propagated count was selected, with Cell Ontology identifier as the final deterministic tie-breaker. The selected anchor was replaced by that parent, and every original fine term assigned to the selected anchor was reassigned to the parent. This process was repeated until 30 ontology anchors remained. No ontology nodes were excluded from eligibility as anchors in the analysis reported here.

Algorithm 1 gives a conceptual specification of the reduction. The tuples in the two minimisation steps encode the deterministic tie-breaking rules described above.

The procedure produced 30 ontology-derived labels plus unknown. Because observed frequencies and direct-parent choices drove the ascent, the final anchors could occur at different ontology depths, and one retained anchor could be an ancestor of another; this is therefore a frequency-adaptive ontology reduction, not a fixed-depth or antichain cut. This unequal resolution was intentional: well-supported granular populations could remain separate, whereas less-supported populations were promoted to broader ontology terms. Here, support was relative to the other anchors during reduction at fixed G, rather than an absolute minimum-cell threshold. When both an ancestor and one of its descendants were retained, the ancestor label represented only the residual fine terms assigned to that anchor, rather than the complete ontological extension of the ancestor. For all coarse-label analyses, cells mapped to unknown or the fallback label other were removed. In the validation partition, 748 cells were labelled unknown; consequently, 29,252 cells contributed to coarse-label metrics.

**Algorithm 1.**
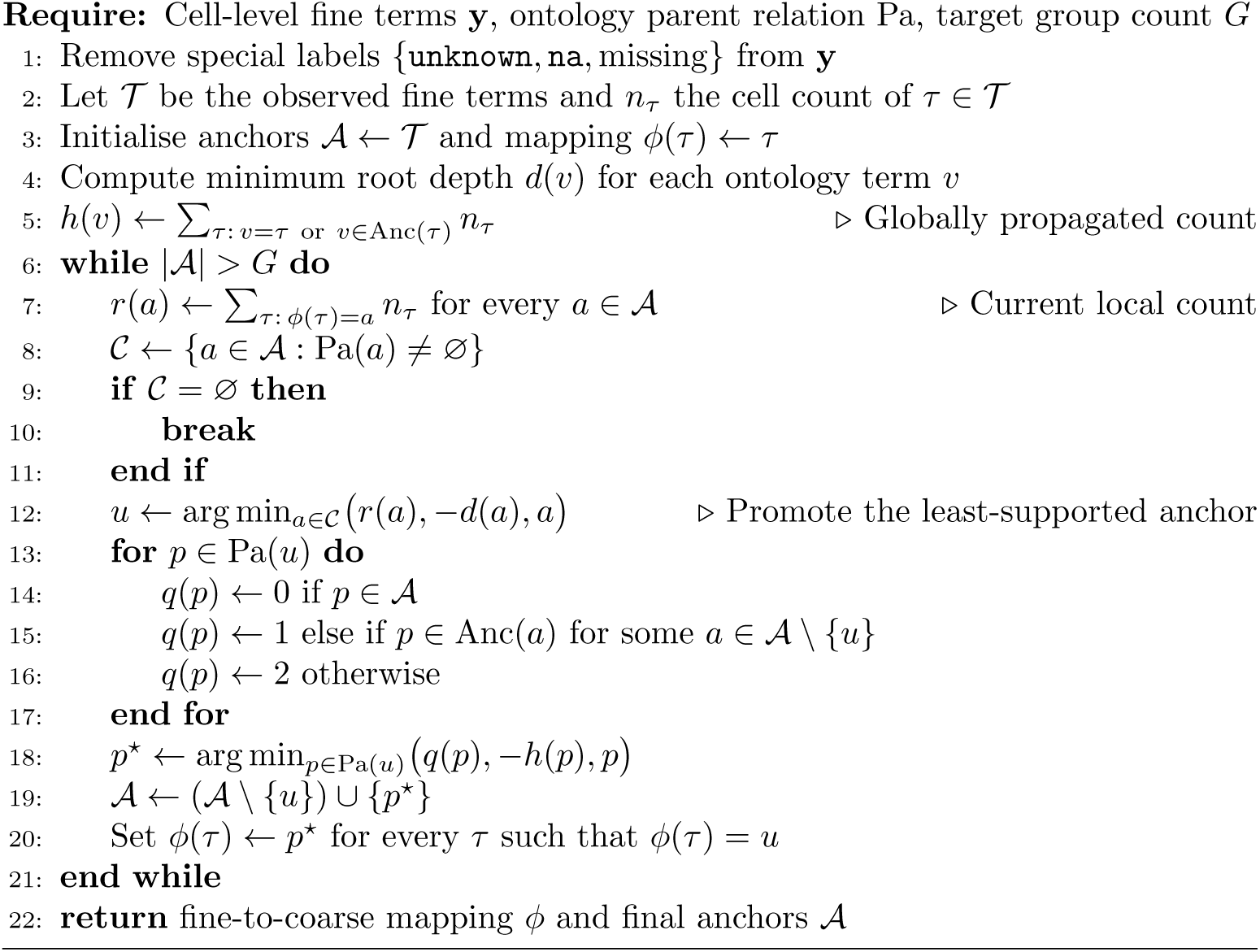
Frequency-adaptive reduction of fine Cell Ontology terms.

### 4.3 Input construction and masking

#### 4.3.1 Ranked Gene Identity formulation

For each cell, expression values were first divided by their sum. To construct the ranking statistic, each gene’s within-cell relative expression was then divided by its mean non-zero relative expression across all filtered Census observations; this statistic was precomputed before the train/validation/test split. Genes with a corpus mean numerically equal to zero were assigned a denominator of one. Expressed genes were ranked in decreasing order of this normalised statistic. When a cell contained more expressed genes than the context length, expressed genes were randomly subsampled to the requested length and then restored to their global expression-rank order. Cells with fewer expressed genes were padded to the context length.

We evaluated nominal context lengths of 128, 256, 512, 1,024, and 2,048 tokens. At each iteration, 15% of non-padding gene tokens were independently selected with uniform probability and replaced by a learned mask token. The model predicted the original gene identity at masked positions using categorical cross-entropy over the 61,497 genes plus special tokens.

The rank of each selected gene in the full within-cell gene ordering was supplied through a sinusoidal positional encoding. Padding positions were assigned no positional signal.

#### 4.3.2 Binned Gene Expression formulation

For each cell, zero expression values were retained as bin 0. Non-zero values were discretised by comparison with 50 within-cell quantiles, producing integer expression targets from 1 to 50. A context contained approximately (1 − m/2)S expressed genes (when available) and (m/2)S unexpressed genes, where S is the context length and m = 0.15 is the masking probability. The two requested counts were independently rounded down, giving effective sequence lengths of 127, 255, 511, 1,023, and 2,047 positions for nominal context lengths of 128, 256, 512, 1,024, and 2,048, respectively. Expressed and unexpressed candidates were randomly sampled when more candidates were available than required. Each selected position was subsequently masked independently with probability 0.15 by replacing its expression input with −1; gene identifiers remained visible.

The binned model used two prediction heads. An expression head was optimised by mean-squared error on positions that were both masked and non-zero in the target,

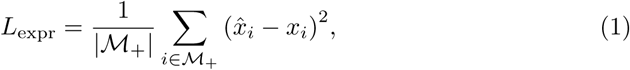

where M_+_ = {i : i is masked and x*_i_* > 0}. A second head predicted whether each masked gene was active using binary cross-entropy, L_act_. Training minimised

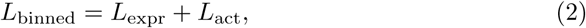

with equal weights. Scaling plots for this formulation report L_expr_, not the summed optimisation objective.

The validation partition contained the same cell identities at every evaluation, but the dataset pipeline recomputed stochastic gene selection and masking at each validation event. Validation losses therefore used newly sampled masks and gene subsets. For representation evaluation, embeddings were recomputed from the corresponding selected but unmasked inputs, so their variation across evaluations reflected gene selection rather than masking.

### 4.4 Transformer architecture

Both formulations used a custom encoder-only [27] transformer [28]. Each pre-normalised block consisted of four-head scaled dot-product self-attention followed by a feed-forward transition with hidden dimension 4W, ReLU activation, and dropout. Dropout with probability 0.2 was applied to the attention and feed-forward outputs before their residual additions and between the two feed-forward linear layers. The ranked formulation additionally applied dropout with probability 0.2 after adding the sinusoidal rank encoding, and each prediction head applied dropout between its two linear layers. No dropout was applied to the attention weights themselves. Query, key, and value projections had no bias; the attention output projection used a bias. The attention gate available in the software was disabled.

In the ranked formulation, gene identifiers were mapped to learned W -dimensional embeddings, combined with sinusoidal rank encodings, and decoded by a two-layer gene classifier with ReLU and dropout. In the binned formulation, learned gene embeddings and a two-layer projection of the scalar expression bin were separately layer-normalised and summed. Separate two-layer decoders produced the expression-bin estimate and activity logit.

We use N to denote the trainable non-embedding parameter count. Thus N excludes the gene/expression input modules and all task-specific decoders. The total trainable parameter count, used only in explicitly labelled supplementary diagnostics, is denoted by N_tot_. Encoder depth, D, is the number of transformer blocks; encoder width, W, is the hidden dimension. The architectural shape variable R, or depth-to-width ratio, was defined as

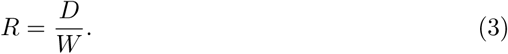

### 4.5 Optimisation and training

Models were initialised with AdamW [29] using β_1_ = 0.9, β_2_ = 0.95, PyTorch’s default ɛ = 10*^−^*^8^, and weight decay 0.01 applied through a single parameter group. For one-cycle [30] runs, PyTorch’s default momentum cycling varied AdamW’s β_1_ from 0.95 to 0.85 during the increasing-learning-rate phase and back to 0.95 during annealing, while β_2_ remained 0.95. The one-cycle schedule used a 30% increasing phase, cosine annealing, initial learning rate λ_max_/25, and final learning rate λ_max_/250,000. The warm-up/constant sweeps did not cycle momentum and retained the initialised AdamW coefficients. The gradient norm was clipped to one [31]. Training used bfloat16 [32] mixed precision, a physical and effective batch size of 32 cells, and no gradient accumulation.

#### 4.5.1 Batch-size selection

Before the principal sweeps, we evaluated physical batch sizes of 4, 16, 32, 64, and 128 for both formulations. These experiments used context length 256, six transformer blocks, four attention heads, widths of 128, 256, 512, and 1,024, learning rate 3 × 10*^−^*^4^, a one-cycle schedule, and 15,000 optimiser steps. Batch size 32 was selected for subsequent experiments because it was the smallest tested value comfortably above the low-batch regime while retaining modest memory requirements.

#### 4.5.2 Context-length and non-embedding-parameter sweep

The context-length/model-size sweep used a one-cycle learning-rate schedule with maximum learning rate 10*^−^*^3^ and up to 50,000 optimiser steps. It crossed five context lengths (128–2,048), four widths (128, 256, 512, and 768), and five depths (2–6), for 100 planned configurations per formulation. The analysed data contain 99 ranked and 100 binned configurations. A complete 50,000-step run with a batch size of 32 produced 1.6 million training-cell examples, and training stopped at the optimiser-step limit before exhausting the training partition. Validation and downstream representation evaluation were performed before the first optimiser update and every 500 optimiser steps thereafter.

#### 4.5.3 Compute-matched sweeps

Learning-rate and depth/width sweeps used a linear warm-up from 10*^−^*^6^ times the target learning rate to the target value over 500 optimiser steps, followed by a constant learning rate. Because configurations differed in FLOPs per step, the maximum step count was selected separately for each architecture to approach its target compute budget. Runs that reached at least 90% of the requested slice were eligible for learning-rate optimisation; response-surface observations were retained when they reached at least 99% of their requested compute slice.

### 4.6 Parameter and compute accounting

Parameter counts were obtained directly from the instantiated PyTorch modules. Forward-pass floating-point operations per cell, F_fwd_, were estimated with Lightning Fabric’s operator-level FLOP counter on a meta-device execution of the complete model forward pass. Estimated training FLOPs at optimiser step t were

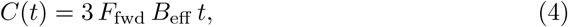

where B_eff_ is the effective batch size. The factor of three approximates one forward and two backward-equivalent passes. This measure includes model-forward operators but is not an end-to-end hardware measurement; it excludes data loading, optimiser updates, and device utilisation overhead. The FLOP counter used the nominal configured context length, including for the binned formulation, and therefore slightly overestimates the operations associated with its effective S − 1-position inputs.

### 4.7 Scaling with non-embedding parameter count and context length

For the initial scaling analysis, models were compared at the largest optimiser step reached by every run within a formulation. This was step 37,499 for the ranked formulation and step 49,999 for the binned formulation. At each context length, we fitted

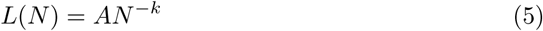

by ordinary least squares after base-10 logarithmic transformation of L and N. We neglected an irreducible component of the loss [33], as our purpose was to investigate local, rather than asymptotic, behaviour. The largest-N configuration was excluded from the fit and used as a held-out extrapolation target. The reported exponent is k = −b_size_, where b_size_ is the fitted slope of log_10_ L on log_10_ N. These comparisons hold optimiser steps, batch size, and the number of examples seen approximately fixed, but not training FLOPs; larger encoders require more compute per step.

### 4.8 Learning-rate scaling

Learning-rate sweeps crossed four target non-embedding parameter counts (10^6^, 10^7^, 10^8^, 10^9^), three target depths (2, 6, and 20 blocks), and five learning rates (10*^−^*^5^, 3 × 10*^−^*^5^, 10*^−^*^4^, 3 × 10*^−^*^4^, 10*^−^*^3^). Width was chosen in multiples of four to minimise the absolute difference between the actual and target non-embedding parameter count while retaining four attention heads. The ranked and binned sweeps targeted total budgets of 7.0 × 10^17^ and 1.4 × 10^17^ estimated training FLOPs, respectively.

Loss was sampled at eight equally spaced fractions from 9.375% to 75% of each matched budget. Thus the penultimate slice shown in Figure 4 corresponds to 65.625% of the target budget: 4.59 × 10^17^ FLOPs for the ranked formulation and 9.19 × 10^16^ FLOPs for the binned formulation.

For each non-embedding-parameter/depth pair at a given compute slice, we first selected the sampled learning rate with the smallest loss. If that point had immediate sampled neighbours on both sides in the original learning-rate grid, and all three runs had finite losses and reached the completion threshold, we fitted

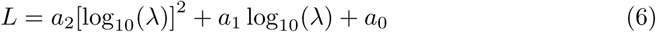

to the best point and its two neighbours. We accepted the continuous optimum log_10_ λ*^∗^* = −a_1_/(2a_2_) only when a_2_ > 0 and the vertex lay inside the three-point fitting interval. Otherwise, including when the discrete optimum occurred at a sweep boundary, or either immediate neighbour was incomplete, missing, or non-finite, we retained the sampled optimum.

We compared four descriptive scaling rules,

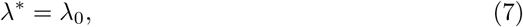

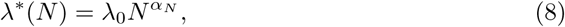

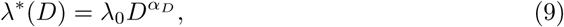

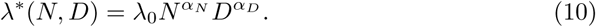

The models were fitted by ordinary least squares in base-10 log space. Fit quality was summarised by root mean square error in log_10_ λ (dex). These errors describe the observed architecture grid and are not out-of-distribution test errors. For coefficient sensitivity intervals, architecture points were resampled with replacement 2,000 times and the joint model was refitted; percentile intervals were obtained from the bootstrap distribution.

### 4.9 Depth-to-width response surfaces

The architectural-shape sweep crossed seven target non-embedding parameter counts (1 million, 3 million, 10 million, 30 million, 100 million, 300 million, and 1 billion) with seven target depths (2, 4, 8, 16, 24, 32, and 64 blocks), yielding 49 configurations per formulation. Width was selected to match each target size as closely as possible, while the number of attention heads was fixed at 4. Learning rates were assigned using the joint size/depth rule estimated in the preceding sweep. Each run targeted up to 10^18^ estimated training FLOPs, subject to a maximum of 100,000 optimiser steps.

Training curves were sampled at seven logarithmically spaced compute budgets between 3 × 10^15^ and 10^18^ FLOPs. Observations with completion fraction below 0.99 at a requested slice were excluded. Let

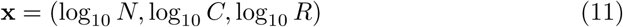

and let z*_i_* = (x*_i_* − x̄*_i_*)/s*_i_* denote the standardised predictors over the retained observations. We fitted the raw pre-training loss by ordinary least squares using the full quadratic model

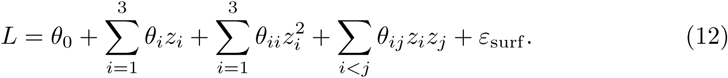

Predictive performance was evaluated with random five-fold cross-validation (seed 7) and leave-one-compute-slice-out validation. The former splits surface observations and may therefore place different compute slices from the same training run in training and test folds; the latter evaluates interpolation to an unseen compute slice while retaining the same architecture grid.

For fixed N and C, the optimal log depth-to-width ratio was the analytic stationary point along the z*_R_* axis. An optimum was accepted only when the fitted coefficient of *z*^2^_*R*_ was positive. The optimum was unconstrained and could fall outside the observed range of ratios; such cases were marked as extrapolations.

Uncertainty bands were generated from 250 row-bootstrap samples (seed 11). Each sample refitted the response surface to observations drawn with replacement, and non-convex fits in the log-ratio direction were discarded. Curves show the 5th and 95th percentiles of the retained optima. Because compute slices from a single run are correlated, these bands should be interpreted as surface-sensitivity intervals rather than independent-run confidence intervals.

### 4.10 Downstream representation evaluation

At validation checkpoints, models were placed in evaluation mode,, and token-level encoder outputs from the selected but unmasked inputs were averaged across sequence positions to obtain a single vector per cell. Embedding inference followed the training mixed-precision configuration, and the saved embeddings were converted to 32-bit floating-point before downstream analysis. No batch integration or correction algorithm was applied. The same held-out validation cells used for pre-training validation were used for the representation analyses below; downstream labels were not used to update the encoder.

For clustering, cell embeddings were placed in AnnData.X and passed to Scanpy with its representation argument left at the default. Because all analysed embeddings had more than 50 dimensions, Scanpy projected them to 50 principal components before constructing a 15-nearest-neighbour graph with Euclidean distance. Leiden clustering was then performed at fixed resolution 1.0. Fine-label clustering used all 30,000 validation cells. For coarse-label clustering, cells without a valid coarse label were first removed, and the PCA, neighbour graph, and Leiden clustering were recomputed on the remaining 29,252 cells. Agreement with the corresponding labels was quantified with normalised mutual information (NMI), adjusted Rand index (ARI), and homogeneity.

For the linear-probe analysis, cells were divided into stratified 80% training and 20% validation subsets with seed 0. A closed-form ridge-regression classifier without an intercept was fitted to one-hot targets using penalty ρ = 1,

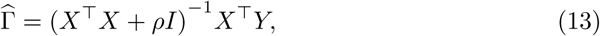

implemented through a least-squares solve. The mean-pooled embeddings were used without centring, feature standardisation, dimensionality reduction, or length normalisation. Accuracy was reported on the 20% probe-validation subset for both fine and coarse labels.

We additionally calculated scIB [34] metrics on the uncorrected embeddings using the source dataset as the batch label and cell type as the biological label. These calculations used the raw, unprojected mean-pooled embeddings. For the scIB NMI and ARI, the Leiden resolution was selected by maximising fine-label NMI across 10 resolutions from 0.2 to 2.0, following the scIB default procedure. Label average silhouette width (ASW) was linearly scaled to the interval [0, 1]. Batch ASW was calculated within each fine cell type and averaged across eligible cell types, with larger values indicating greater mixing of source datasets. Graph connectivity was the mean across fine cell types of the fraction of cells in the largest connected component of that cell type’s neighbour subgraph. Because no integration method was applied, these values characterise biological preservation and batch structure in the learned representation rather than the performance of a batch-correction algorithm.

For compact visualisation, we defined a study-specific composite score as the unweighted row mean of the available values among fine NMI, fine ARI, fine homogeneity, coarse NMI, coarse ARI, coarse homogeneity, label ASW, graph connectivity, and coarse-label ridge validation accuracy:

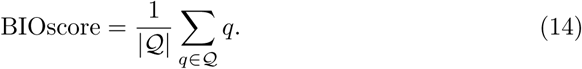

All component metrics were oriented so that larger values indicate better performance. The fine-and coarse-NMI, ARI, and homogeneity terms in BIOscore were the fixed-resolution clustering metrics described above, rather than the resolution-optimised scIB NMI and ARI. Neither scIB NMI, scIB ARI, nor batch ASW entered BIOscore. BIOscore is not a standard scIB statistic; component metrics are reported separately to avoid relying solely on the composite.

### 4.11 Software and reproducibility

The pipeline was implemented in Python 3.12 using PyTorch, Lightning [35], TileDB-SOMA, TileDB-SOMA-ML, Polars, Scanpy [36], scIB, scikit-learn [37], and Aim. The locked GPU environment used PyTorch 2.11.0+cu128, Lightning 2.6.1, TileDB-SOMA 2.3.0, TileDB-SOMA-ML 0.1.0, Scanpy 1.11.5, and scIB 1.1.7. The training entry point set PyTorch’s float32 matrix-multiplication precision to medium. Each training task used one accelerator: either an NVIDIA RTX PRO 6000 Blackwell GPU with 96 GB of memory on an on-premises cluster or a custom NVIDIA Ampere A100 GPU with 64 GB of HBM2e memory on CINECA’s LEONARDO system.

The code will be made available as a GitHub repository upon publication.

## Acknowledgements

We thank Michelangelo Nardi for helping in running some preliminary experiments that were not included in this work.

## Notes

### Competing Interest Statement

The authors have declared no competing interest.

## References

[1] Hestness, J., Narang, S., Ardalani, N., Diamos, G.F., Jun, H., Kianinejad, H., Patwary, M.M.A., Yang, Y., Zhou, Y.: Deep learning scaling is predictable, empirically. CoRR abs/1712.00409 (2017) 1712.00409

[2] Kaplan, J., McCandlish, S., Henighan, T., Brown, T.B., Chess, B., Child, R., Gray, S., Radford, A., Wu, J., Amodei, D.: Scaling laws for neural language models. CoRR abs/2001.08361 (2020) 2001.08361

[3] Hoffmann, J., Borgeaud, S., Mensch, A., Buchatskaya, E., Cai, T., Rutherford, E., Las Casas, D., Hendricks, L.A., Welbl, J., Clark, A., Hennigan, T., Noland, E., Millican, K., Driessche, G., Damoc, B., Guy, A., Osindero, S., Simonyan, K., Elsen, E., Rae, J.W., Vinyals, O., Sifre, L.: Training Compute-Optimal Large Language Models (2022). https://arxiv.org/abs/2203.15556

[4] Regev, A., Teichmann, S.A., Lander, E.S., Amit, I., Benoist, C., Birney, E., Bodenmiller, B., Campbell, P., Carninci, P., Clatworthy, M., Clevers, H., Deplancke, B., Dunham, I., Eberwine, J., Eils, R., Enard, W., Farmer, A., Fugger, L., Göttgens, B., Hacohen, N., Haniffa, M., Hemberg, M., Kim, S., Klenerman, P., Kriegstein, A., Lein, E., Linnarsson, S., Lundberg, E., Lunde-berg, J., Majumder, P., Marioni, J.C., Merad, M., Mhlanga, M., Nawijn, M., Netea, M., Nolan, G., Pe’er, D., Phillipakis, A., Ponting, C.P., Quake, S., Reik, W., Rozenblatt-Rosen, O., Sanes, J., Satija, R., Schumacher, T.N., Shalek, A., Shapiro, E., Sharma, P., Shin, J.W., Stegle, O., Stratton, M., Stubbington, M.J.T., Theis, F.J., Uhlen, M., Oudenaarden, A., Wagner, A., Watt, F., Weissman, J., Wold, B., Xavier, R., Yosef, N.a.: The human cell atlas. eLife 6 (2017) 10.7554/elife.27041

[5] Theodoris, C.V., Xiao, L., Chopra, A., Chaffin, M.D., Al Sayed, Z.R., Hill, M.C., Mantineo, H., Brydon, E.M., Zeng, Z., Liu, X.S., Ellinor, P.T.: Transfer learning enables predictions in network biology. Nature 618(7965), 616–624 (2023) 10.1038/s41586-023-06139-9

[6] Cui, H., Wang, C., Maan, H., Pang, K., Luo, F., Duan, N., Wang, B.: scgpt: toward building a foundation model for single-cell multi-omics using generative ai. Nature Methods 21(8), 1470–1480 (2024) 10.1038/s41592-024-02201-0

[7] Brown, T.B., Mann, B., Ryder, N., Subbiah, M., Kaplan, J., Dhariwal, P., Neelakantan, A., Shyam, P., Sastry, G., Askell, A., Agarwal, S., Herbert-Voss, A., Krueger, G., Henighan, T., Child, R., Ramesh, A., Ziegler, D.M., Wu, J., Winter, C., Hesse, C., Chen, M., Sigler, E., Litwin, M., Gray, S., Chess, B., Clark, J., Berner, C., McCandlish, S., Radford, A., Sutskever, I., Amodei, D.: Language Models are Few-Shot Learners (2020). https://arxiv.org/abs/2005.14165

[8] Hao, M., Gong, J., Zeng, X., Liu, C., Guo, Y., Cheng, X., Wang, T., Ma, J., Zhang, X., Song, L.: Large-scale foundation model on single-cell transcriptomics. Nature Methods 21(8), 1481–1491 (2024) 10.1038/s41592-024-02305-7

[9] Fischer, F., Fischer, D.S., Mukhin, R., Isaev, A., Biederstedt, E., Villani, A.-C., Theis, F.J.: sctab: Scaling cross-tissue single-cell annotation models. Nature Communications 15(1) (2024) 10.1038/s41467-024-51059-5

[10] Rizvi, S.A., Levine, D., Patel, A., Zhang, S., Wang, E., Perry, C.J., Vrkic, I., Constante, N.M., Fu, Z., He, S., Zhang, D., Tang, C., Lyu, Z., Darji, R., Li, C., Sun, E., Jeong, D., Zhao, L., Kwan, J., Braun, D., Hafler, B., Chung, H., Dhodapkar, R.M., Jaeger, P., Perozzi, B., Ishizuka, J., Azizi, S., Dijk, D.: Scaling large language models for next-generation single-cell analysis (2025) 10.1101/2025.04.14.648850

[11] Adduri, A.K., Gautam, D., Bevilacqua, B., Imran, A., Shah, R., Naghipourfar, M., Teyssier, N., Ilango, R., Nagaraj, S., Dong, M., Ricci-Tam, C., Carpenter, C., Subramanyam, V., Winters, A., Tirukkovular, S., Sullivan, J., Plosky, B.S., Eraslan, B., Youngblut, N.D., Leskovec, J., Gilbert, L.A., Konermann, S., Hsu, P.D., Dobin, A., Burke, D.P., Goodarzi, H., Roohani, Y.H.: Predicting cellular responses to perturbation across diverse contexts with state. bioRxiv (2025) 10.1101/2025.06.26.661135 https://www.biorxiv.org/content/early/2025/07/10/2025.06.26.661135.full.pdf

[12] Gandhi, S., Javadi, F., Svensson, V., Khan, U., Jones, M.G., Yu, J., Merico, D., Goodarzi, H., Alidoust, N.: Tahoe-x1: Scaling perturbation-trained single-cell foundation models to 3 billion parameters. bioRxiv (2025) 10.1101/2025.10.23.683759 https://www.biorxiv.org/content/early/2025/10/23/2025.10.23.683759.full.pdf

[13] Dong, M., Adduri, A., Gautam, D., Carpenter, C., Shah, R., Ricci-Tam, C., Kluger, Y., Burke, D.P., Roohani, Y.H.: Stack: In-context learning of single-cell biology. bioRxiv (2026) 10.64898/2026.01.09.698608

[14] Chen, H., Venkatesh, M.S., Gómez Ortega, J., Mahesh, S.V., Nandi, T.N., Madduri, R.K., Pelka, K., Theodoris, C.V.: Scaling and quantization of large-scale foundation model enables resource-efficient predictions in network biology. Nature Computational Science 6(5), 450–463 (2026) 10.1038/s43588-026-00972-4

[15] DenAdel, A., Hughes, M., Thoutam, A., Gupta, A., Navia, A.W., Fusi, N., Raghavan, S., Winter, P.S., Amini, A.P., Crawford, L.: Evaluating the role of pretraining dataset size and diversity on single-cell foundation model performance. Nature Methods 23(7), 1447–1457 (2026) 10.1038/s41592-026-03120-y

[16] Program, C.S.-C.B., Abdulla, S., Aevermann, B., Assis, P., Badajoz, S., Bell, S.M., Bezzi, E., Cakir, B., Chaffer, J., Chambers, S., Michael Cherry, J., Chi, T., Chien, J., Dorman, L., Garcia-Nieto, P., Gloria, N., Hastie, M., Hegeman, D., Hilton, J., Huang, T., Infeld, A., Istrate, A.-M., Jelic, I., Katsuya, K., Kim, Y.J., Liang, K., Lin, M., Lombardo, M., Marshall, B., Martin, B., McDade, F., Megill, C., Patel, N., Predeus, A., Raymor, B., Robatmili, B., Rogers, D., Rutherford, E., Sadgat, D., Shin, A., Small, C., Smith, T., Sridharan, P., Tarashansky, A., Tavares, N., Thomas, H., Tolopko, A., Urisko, M., Yan, J., Yeretssian, G., Zamanian, J., Mani, A., Cool, J., Carr, A.: Cz cell×gene discover: A single-cell data platform for scalable exploration, analysis and modeling of aggregated data. bioRxiv (2023) 10.1101/2023.10.30.563174 https://www.biorxiv.org/content/early/2023/11/02/2023.10.30.563174.full.pdf

[17] Kedzierska, K.Z., Crawford, L., Amini, A.P., Lu, A.X.: Zero-shot evaluation reveals limitations of single-cell foundation models. Genome Biology 26(1) (2025) 10.1186/s13059-025-03574-x

[18] Ahlmann-Eltze, C., Huber, W., Anders, S.: Deep-learning-based gene perturbation effect prediction does not yet outperform simple linear baselines. Nature Methods 22(8), 1657–1661 (2025) 10.1038/s41592-025-02772-6

[19] Yang, G., Yu, D., Zhu, C., Hayou, S.: Tensor programs VI: Feature learning in infinite depth neural networks. In: The Twelfth International Conference on Learning Representations (2024). https://openreview.net/forum?id=17pVDnpwwl

[20] Ren, L., Liu, Y., Shen, Y., Chen, W.: Rethinking Language Model Scaling under Transferable Hypersphere Optimization (2026). https://arxiv.org/abs/2603.28743

[21] Abnar, S., Shah, H., Busbridge, D., El-Nouby, A., Susskind, J.M., Thilak, V.: Parameters vs FLOPs: Scaling laws for optimal sparsity for mixture-of-experts language models. In: Singh, A., Fazel, M., Hsu, D., Lacoste-Julien, S., Berkenkamp, F., Maharaj, T., Wagstaff, K., Zhu, J. (eds.) Proceedings of the 42nd International Conference on Machine Learning. Proceedings of Machine Learning Research, vol. 267, pp. 204–230. PMLR,(2025). https://proceedings.mlr.press/v267/abnar25a.html

[22] Yang, G., Hu, E.J., Babuschkin, I., Sidor, S., Liu, X., Farhi, D., Ryder, N., Pachocki, J., Chen, W., Gao, J.: Tensor Programs V: Tuning Large Neural Networks via Zero-Shot Hyperparameter Transfer (2022). https://arxiv.org/abs/2203.03466

[23] Mlodozeniec, B.K., Ablin, P., Béthune, L., Busbridge, D., Klein, M., Ramapuram, J., Cuturi, M.: Completed hyperparameter transfer across modules, width, depth, batch and duration. In: The Fourteenth International Conference on Learning Representations (2026). https://openreview.net/forum?id=elB9k4nTL1

[24] Paszke, A., Gross, S., Massa, F., Lerer, A., Bradbury, J., Chanan, G., Killeen, T., Lin, Z., Gimelshein, N., Antiga, L., Desmaison, A., Köpf, A., Yang, E., DeVito, Z., Raison, M., Tejani, A., Chilamkurthy, S., Steiner, B., Fang, L., Bai, J., Chintala, S.: PyTorch: An Imperative Style, High-Performance Deep Learning Library (2019). https://arxiv.org/abs/1912.01703

[25] Harris, C.R., Millman, K.J., Walt, S.J., Gommers, R., Virtanen, P., Cournapeau, D., Wieser, E., Taylor, J., Berg, S., Smith, N.J., Kern, R., Picus, M., Hoyer, S., Kerkwijk, M.H., Brett, M., Haldane, A., Río, J.F., Wiebe, M., Peterson, P., Gérard-Marchant, P., Sheppard, K., Reddy, T., Weckesser, W., Abbasi, H., Gohlke, C., Oliphant, T.E.: Array programming with NumPy. Nature 585(7825), 357–362 (2020) 10.1038/s41586-020-2649-2

[26] Bard, J., Rhee, S.Y., Ashburner, M.: An ontology for cell types. Genome Biology 6(2) (2005) 10.1186/gb-2005-6-2-r21

[27] Devlin, J., Chang, M.-W., Lee, K., Toutanova, K.: BERT: Pre-training of deep bidirectional transformers for language understanding. In: Burstein, J., Doran, C., Solorio, T. (eds.) Proceedings of the 2019 Conference of the North American Chapter of the Association for Computational Linguistics: Human Language Technologies, Volume 1 (Long and Short Papers), pp. 4171–4186. Association for Computational Linguistics, Minneapolis, Minnesota (2019). 10.18653/v1/N19-1423. https://aclanthology.org/N19-1423/

[28] Vaswani, A., Shazeer, N., Parmar, N., Uszkoreit, J., Jones, L., Gomez, A.N., Kaiser, L., Polosukhin, I.: Attention is all you need. In: Proceedings of the 31st International Conference on Neural Information Processing Systems. NIPS’17, pp. 6000–6010. Curran Associates Inc., Red Hook, NY, USA (2017)

[29] Loshchilov, I., Hutter, F.: Decoupled weight decay regularization. In: International Conference on Learning Representations (2019). https://openreview.net/forum?id=Bkg6RiCqY7

[30] Smith, L.N., Topin, N.: Super-Convergence: Very Fast Training of Neural Networks Using Large Learning Rates (2018). https://arxiv.org/abs/1708.07120

[31] Pascanu, R., Mikolov, T., Bengio, Y.: On the difficulty of training recurrent neural networks. In: Proceedings of the 30th International Conference on International Conference on Machine Learning -Volume 28. ICML’13, pp. 1310–1318. JMLR.org,(2013)

[32] Kalamkar, D., Mudigere, D., Mellempudi, N., Das, D., Banerjee, K., Avancha, S., Vooturi, D.T., Jammalamadaka, N., Huang, J., Yuen, H., Yang, J., Park, J., Heinecke, A., Georganas, E., Srinivasan, S., Kundu, A., Smelyanskiy, M., Kaul, B., Dubey, P.: A Study of BFLOAT16 for Deep Learning Training (2019). https://arxiv.org/abs/1905.12322

[33] Kendiukhov, I.: Scaling laws for masked-reconstruction transformers on single-cell transcriptomics. Transactions on Machine Learning Research (2026)

[34] Luecken, M.D., Büttner, M., Chaichoompu, K., Danese, A., Interlandi, M., Mueller, M.F., Strobl, D.C., Zappia, L., Dugas, M., Colomé-Tatché, M., Theis, F.J.: Benchmarking atlas-level data integration in single-cell genomics. Nature Methods 19(1), 41–50 (2021) 10.1038/s41592-021-01336-8

[35] Falcon, W., Borovec, J., Wälchli, A., Eggert, N., Schock, J., Jordan, J., Skafte, N., Bereznyuk, V., Harris, E., Tullie Murrell, Yu, P., Præsius, S., Addair, T., Zhong, J., Lipin, D., Uchida, S. Shreyas Bapat, Schröter, H., Dayma, B., Karnachev, A., Akshay Kulkarni, Shunta Komatsu, Martin B, Jean-Baptiste Schiratti, Mary, H., Byrne, D., Cristobal Eyzaguirre, Cinjon, Bakhtin, A.: PyTorchLightning/pytorch-lightning: 0.7.6 release. Zenodo (2020). 10.5281/ZENODO.3828935. https://zenodo.org/record/3828935

[36] Wolf, F.A., Angerer, P., Theis, F.J.: Scanpy: large-scale single-cell gene expression data analysis. Genome Biology 19(1) (2018) 10.1186/s13059-017-1382-0

[37] Pedregosa, F., Varoquaux, G., Gramfort, A., Michel, V., Thirion, B., Grisel, O., Blondel, M., Prettenhofer, P., Weiss, R., Dubourg, V., Vanderplas, J., Passos, A., Cournapeau, D., Brucher, M., Perrot, M., Duchesnay, E.: Scikit-learn: Machine learning in Python. Journal of Machine Learning Research 12, 2825–2830 (2011)

